# Chiral mismatch in collagen-mimetic peptides modulates cell migration through integrin-mediated molecular recognition

**DOI:** 10.1101/2024.07.23.604866

**Authors:** Alexandre Remson, David Dellemme, Marine Luciano, Mathieu Surin, Sylvain Gabriele

## Abstract

Chirality influences essential biological processes such as molecular recognition and self-organization, impacting cell proliferation and differentiation mediated by interactions with the extracellular matrix (ECM). Despite extensive research on cell migration, the role of chirality on matrix-cell interactions has been largely overlooked. To explore this aspect, we engineered culture surfaces coated with natural collagen I or collagen-mimetic peptides (CMPs) with opposite chirality, i.e. with either L- or D-amino acids in their sequences. Here we show that D-surfaces impede epithelial keratocyte spreading, making cells more rounded, less adhesive, and slower. Further investigation through integrin inhibition assays and molecular dynamics simulations revealed that a chiral mismatch destabilizes the triple helix of heterochiral CMPs at the L/D junction. This study underscores the profound impact of ECM chirality on cellular behavior, providing new insights into the relationship between matrix (supra)molecular chirality, integrin-mediated molecular recognition, and cell migration dynamics.

---

Chirality is a fundamental characteristic of life^1^, manifesting in the structure of biological molecules, which typically exhibit homochiral units, as observed in DNA and proteins. This raises intriguing questions about why nature predominantly selects left-handed (L) amino acids and right-handed (D) sugars in living organisms. Recent studies have provided insights into the mechanisms of chirality selection, demonstrating diastereoselectivity in prebiotic ligation reactions and the emergence of homochirality through crystallization processes^2^. Numerous biological processes, especially in early development, are influenced by chirality^3,4^. One prominent example is the left-right (L/R) asymmetry in internal organs, where structures like the heart undergo asymmetric rotation to establish functional orientation^5^. Significant changes in Drosophila movements occur with altered expression of the conserved myosin-1D (Myo-1D), underlining how molecular-level chirality impacts the entire organism^6^. On 2D circular fibronectin micropatterns, fibroblasts exhibited chiral arrangements due to actin assembly at peripheral focal adhesions^7^. In 3D environments, cells display spontaneous rotational movements influenced by the ECM components, mediated by proteins like α-actinin-1^8^. At the molecular level, chirality is crucial for the specificity of recognition mechanisms that underlie fundamental cellular processes^9^, such as proliferation^10^ and differentiation^11^. Despite these findings, the impact of ECM chirality on cellular migration remains largely unexplored.

To address this gap, we engineered culture surfaces coated with either natural collagen type I or collagen-mimetic peptides (CMPs) with opposite chiralities (L- and D-amino acids), creating a chiral mismatch that destabilizes the heterochiral peptide at the junction between these amino acids. Our results demonstrate that ECM chirality significantly influences cell morphology and migration. Specifically, D-chiral surfaces hinder epithelial keratocyte spreading, causing cells to become more rounded and reducing their adhesion and migration rates. Further investigation using integrin inhibition assays and molecular dynamics (MD) simulations at the atomic scale revealed that heterochiral peptides presenting right-handed CMPs interact less effectively with the integrin binding site compared to homochiral left-handed CMPs, or natural collagen. MD simulations show that the chiral mismatch at the L/D junction destabilizes the interaction between CMPs and the α1β1 integrin by disorganizing the triple helix structure of the D-peptide. This study highlights the substantial impact of ECM chirality on cellular behavior, offering new perspectives on the intricate relationship between (supra)molecular chirality, integrin-mediated molecular recognition and cell migration dynamics.

## Design and characterization of collagen and CMPs

The effect of ECM chirality on cell migration was investigated using collagen I, representing the major ECM component, and CMPs with (PPG)10 — where P corresponds to proline and G to glycine — in either levorotatory (L) or dextrorotatory (D) stereoisomers **(Extended Fig. 1a)**. Both CMPs are composed of a tetramethylrhodamine fluorescent dye (TAMRA) attached to the N-terminus of the peptide, a single glycine residue acting as a linker, a segment of four lysine (Lys) residues, a segment including two repeats of the tripeptide sequence Proline-Glutamic acid-Glycine (Pro-Glu-Gly), and a segment consisting of ten repeats of the tripeptide sequence Proline-Proline-Glycine (Pro*-Pro*-Gly), where Pro* indicates proline residues with distinct chirality **(Fig. 1a)**. This tripeptide sequence (PPG)10 was shown in previous studies to be a relevant model to mimic collagen, forming a similar PPII triple helix^12^. In our CMPs, the inverse chirality is set on the (PPG)10 sequence only, in order to assess the role of the collagen matrix chirality on cell migration. Hence, the homochiral CMP in **Fig. 1** is the homochiral oligomeric equivalent of natural collagen, whereas the heterochiral CMP possesses a chiral mismatch along the chain between the (PEG)2 section and the (PPG)10 section.

**Figure 1.**
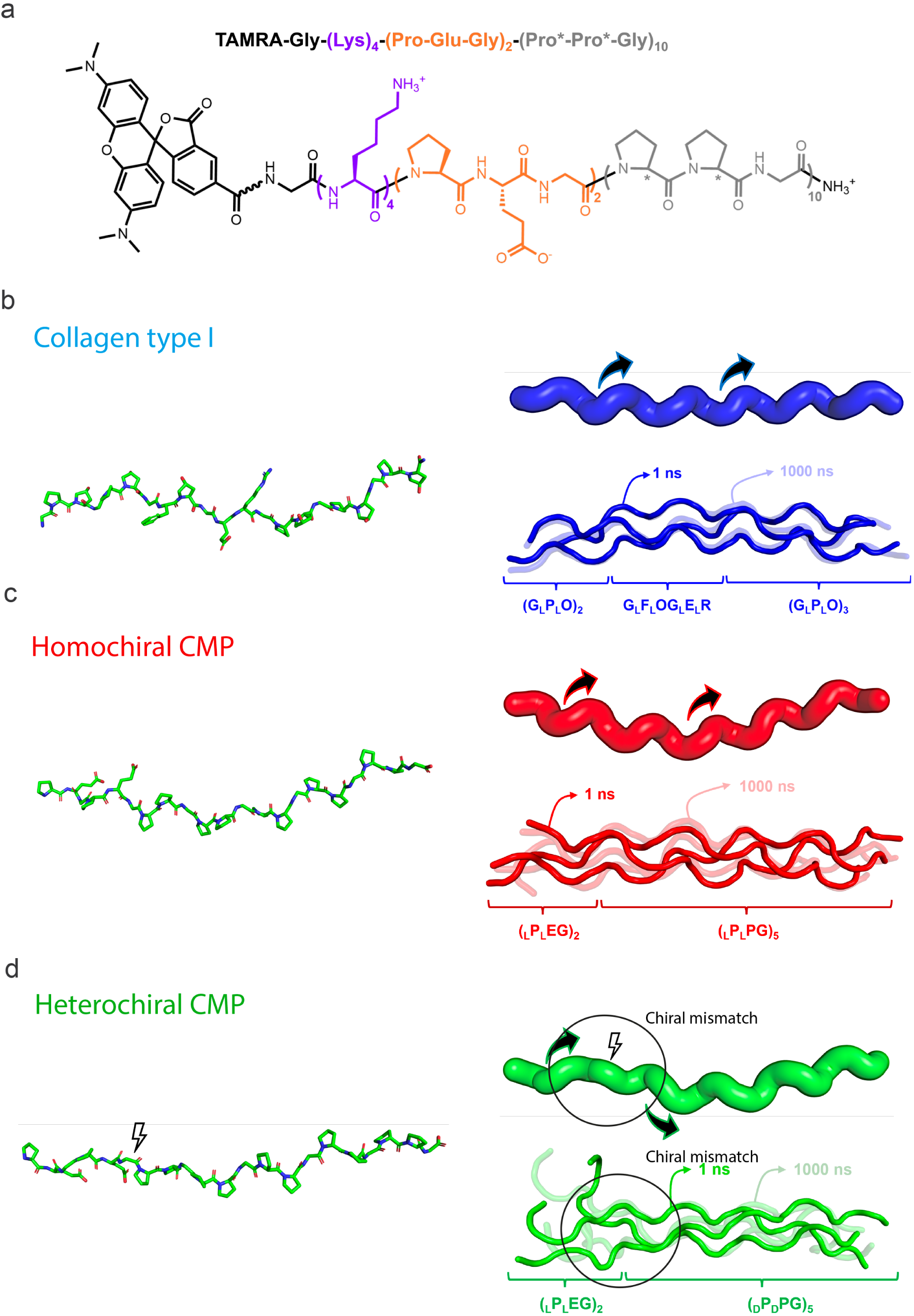
– Collagen mimetic peptides (CMPs) and simplified models for molecular dynamics simulations. **(a)** 2D-representation of the peptidic sequence. From left to right: lysine amino acids in purple, the triplet composed by proline, glutamate and glycine residues in orange, and the proline-rich sequence (PPG) in gray. Simplified models for molecular dynamics (MD) simulations with a shorter peptidic sequence: **(b)** the GFOGER sequence of collagen type-I represented as a single chain and in a triple helix (in blue) after 1 and 1000 11s, and two natural triplets (PEG) following by 5 PPG in both **(c)** natural (homochiral, in red) and **(d)** non-natural (heterochiral, in green) conformations. Visualization of the triple helix resulting from the association of 3 single chains in **(c)** and **(d)** after 1 and 1000 11s. Black circles highlight the chiral mismatch zone on the heterochiral CMP.

Molecular dynamics (MD) simulations in explicit water solvent were performed to investigate the effect of the chiral mismatch on the stability of the triple helix structure of the studied molecules, at the microsecond timescale. In order to reduce the computational cost, simplified models of the experimental peptides were designed with sequences (PEG)2-(PPG)5 **(Fig. 1b-d)**. Let us note that glutamate residues are important for the interaction with integrins^13^ whereas the PPG motif is able to induce the formation of PPII triple helix^14^. In the homochiral peptide, all amino acids are of L-stereochemistry: (LPLEG)2-(LPLPG)5, while in the heterochiral peptide, the prolines of the (PPG)5 part are of D-stereochemistry: (LPLEG)2-(DPDPG)5. These two systems were compared with a model collagen sequence of 21 amino acids, containing the recognition pattern “GFOGER”, identified as an integrin-binding motif^15^. As for collagen, the homochiral peptide is able to form a right-handed supramolecular triple helix, stabilized by hydrogen bonds between the amide groups. Inversely, the PPG residues of the heterochiral peptide, which are of the opposite chirality, induce the formation of a left-handed supramolecular triple helix **(Extended Fig. 1b)**. During the simulations, we observed that the first amino acids of the heterochiral peptide, especially the two “PEG” triplets, were markedly more flexible than their homochiral counterparts, as shown by root-mean-square fluctuation (RMSF) measurements **(Extended Fig. 1c)**. However, the two peptides exhibit a similar low flexibility for the amino acids located in the (PPG)5 triple helix. This suggests that, in the homochiral peptide, the (PEG)2 amino acids are able to maintain the triple helix organization, as the chirality of these units is identical to units in the rest of the strand. In the heterochiral peptide, on the contrary, the L-prolines found in the beginning of the strands are unable to maintain the right helical twist of the D-prolines in the (PPG)5 part, bringing disorder in the supramolecular triple helix **(Fig. 1c and Supplementary Movie 1)**. This trend is reflected by the number of hydrogen bonds inside the triple helix **(Extended Fig. 1d**). The amount of interactions is significantly smaller between the first nine amino acids of the heterochiral peptide (around 4 H-bonds vs 1 H-bond per conformation for the L- and D-peptides, respectively). It reaches the same level for both systems between their last 12 residues (around 6 H-bonds per conformation), which are included in the (PPG)5 region. The model collagen exhibits, in total, slightly more intramolecular H-bonds, which is not surprising as it contains more polar amino acids (hydroxyprolines, GFOGER motif). MD simulations suggest that the chiral mismatch destabilizes the heterochiral peptide at the junction between the L- and D-amino acids, giving more flexibility to the (PEG)2 residues, without influence on the stability of the (PPG)5 triple helix.

We characterized experimentally the natural collagen and the CMPs using circular dichroism (CD) in aqueous solutions. CD spectra exhibited a positive peak at 220 nm for collagen **(Fig. 2a)** and two opposite peaks at 200 nm for L- and D-peptides **(Fig. 2b)** indicating that three (left/right)-polyproline type II helix chains self-assembled together to form a supramolecular triple helix^16^, which is right-handed for collagen I and (LPLPG)10 and left-handed for (DPDPG)10. These assemblies were deposited on flat surfaces (mica or glass) and were characterized by microscopy methods (see Materials & Methods and SI). In the following, we will use the convention “COL” for collagen surfaces, and “L-CMP” for (LPLPG)10 and “D-CMP” for (DPDPG)10 surfaces. Crystallographic parameters enabled to estimate the number of triple helices that self-assembled^12^. We observed that 50 to 600 triples helices self-assembled in length, forming approximately three layers in height. To ensure the preservation of the polyproline type II helix conformation in the three coatings on cell culture substrates, we characterized using circular dichroism (CD) adsorbed layers of COL, L- and D- CMPs on quartz substrates. Culture conditions were mimicked by immersing these substrates overnight in culture medium at room temperature. Characteristic peaks were observed, confirming the conservation of the polyproline type II helix in all coatings **(Fig. 2c).** We then investigated the thermal stability in solution and identified the melting temperatures by CD. To achieve this, the wavelength was set to the polyproline type II helix peaks for COL, L- and D- CMPs **(Fig. 2b)** and the temperature was gradually increased from 10 to 70°C at a rate of 0.25°C/min. Interestingly, the CD signal decreased as the temperature increased **(Fig. 2d)**, suggesting the disruption of hydrogen-bonding interactions between the chains in response to the temperature increase, particularly between the carboxyl group of one chain and the nitrogen group of another chain. Increasing the temperature resulted in a negative/positive (L/D) peak, corresponding to the CD signature of a denatured structure^17^. These spectra were characterized by a sigmoidal behaviour, indicating the cooperativity of the interactions^18^. Cooperativity was evidently higher in COL than in L or D, as the former forms a significantly longer PPII helix than the CMPs, leading to a higher extent of cooperative interactions between the chains^19^. Consequently, the melting temperature was higher (Tm ≈42°C) for collagen I, compared to his homologous L peptide (Tm ≈38°C) and the opposite D peptide (Tm ≈39°C). Given these outcomes, we encountered limitations in utilizing cellular models that required maintenance at 37°C, and opted for a cellular model that is effective at room temperature.

**Figure 2.**
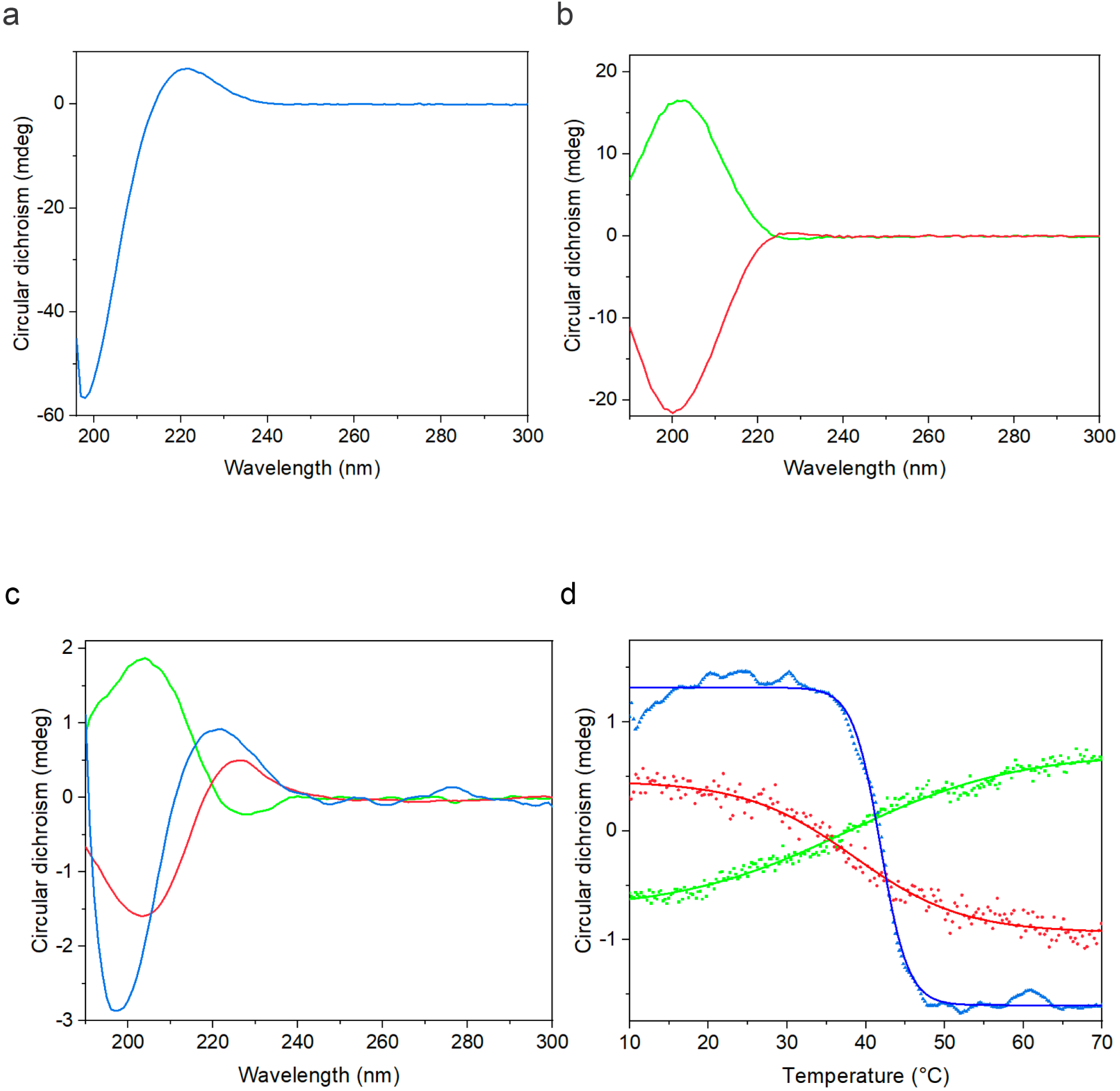
– Circular dichroism (CD) characterization of collagen type I and CMPs. CD spectroscopy **(a)** in Phosphate-Buffered Saline (PBS) for collagen (COL) type I (blue), **(b)** in Milli-Q water of both L- (in red) and D- (in green) CMPs, and **(c)** on quartz surfaces in dry phase for COL (blue), L- (in red) and D- (in green) CMPs. The polyproline type II helix is conserved. **(d)** Thermal analysis of COL (blue), L- (in red) and D- (in green) CMPs at a fixed wavelength (220-225 nm). The temperature was increased slowly by 0.25 deg/min from 10°C to 70°C. Each point corresponds to a CD signal and taken every 0.25°C. The R^2^ of the sigmoidal plots are 0.9726 for COL (in blue), 0.9841 for L- (in red) and 0.9952 for D- (in green) CMPs.

## Matrix chirality modulates epithelial cell morphology

To address this issue, we opted for epithelial keratocytes derived from fish scales, as they can be effectively used at room temperature^20^ **(Fig. 3a)**. Keratocytes offer a unique advantage for the observation of migratory dynamics, given their capability to achieve speeds between 15 and 30 μm/min. Furthermore, they exhibit a distinctive mesenchymal mode of migration, where the movement primarily relies on the polymerization of actin filaments at the leading edge of lamellipodia. This dynamic process propels the plasma membrane forward, coupled with myosin II contraction of cortical actin filaments, resulting in pronounced front-rear polarization^21–25^. Moreoever, keratocytes naturally interface with the internal architecture of fish scales, predominantly composed of collagen^26^. We used microcontact printing (μCP)^27,28^ to create substrates uniformly coated with either COL or CMPs of opposite chirality. The characterization of these substrates using optical microscopy in epifluorescence mode ensured an even distribution and a similar amount of protein or peptides on each surfaces **(Supplementary Fig. 1)**. Fish scales were gently taken off and placed on a glass coverslip to collect an epithelial tissue **(Fig. 3a)**. As reported previously, the internal side of fish scales are composed of collagen^26^ fibers, demonstrating the natural interactions of fish epithelial keratocytes with collagen. Epithelial fish keratocytes attached and spread accross all surfaces, exhibiting distinct morphologies **(Fig. 3b)**. Notably, epithelial cells displayed larger areas on COL, with an average area of 592 ± 150 μm^2^, while the spreading area was 495 ± 150 μm^2^ and 430 ± 130 µm^2^ on L- and D-CMPs, respectively **(Fig. 3c)**. To gain further insight into the impact of chirality changes, we then assessed the cell aspect ratio (CAR). Our results indicated a CAR of ≈2 for collagen and L-surfaces, and ≈1.6 for D-surfaces **(Fig. 3d)**. Interestingly, 3D confocal imaging revealed that the density of actin filaments in the lamellipodia was not equally distributed for the different surfaces. Indeed, the mean gray value was similar on COL and L-CMPs but significantly lower on D-CMPs, suggesting that D-surfaces lead to less branched actin filaments in the lamellipodia which may lead to lower protrusive forces^29^ **(Fig. 3e and Supplementary Fig. 2)**.

**Figure 3.**
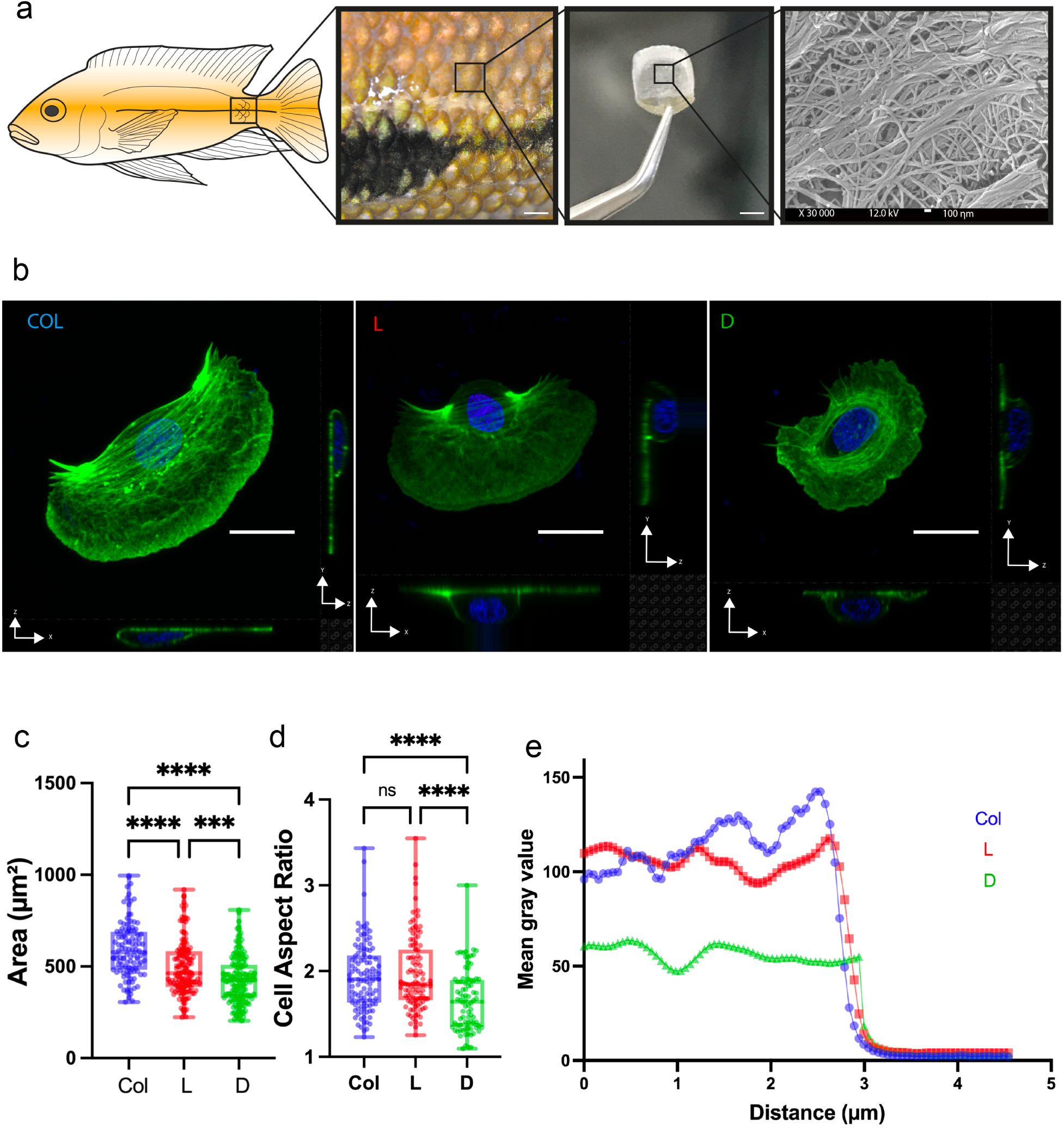
– Matrix chirality modulates cell morphology. **(a)** Representation of a cichlid fish (*Hypsophrys Nicaraguensis*) and the area where scales are extracted. From left to right: black zoom boxes show the visualization of the fish skin, a scale, and the internal architecture of the scale made of collagen as observed by Scanning Electron Microscopy (SEM). From left to right, scale bars are 2 cm, 1 cm and 100 nm. **(b)** Typical confocal images of keratocyte cells on collagen (COL) type I (left), L-CMPs (middle), and D-CMPs (right). Scale bars are 10 µm. **(c)** Spreading area of keratocytes on COL (in blue, n=110), L-CMPs (in red, n=134) and D-CMPs (in green, n=146). **(d)** Cell aspect ratio of keratocytes on COL (in blue, n=110), L- CMPs (in red, n=101) and D-CMPs (in green, n=182). **(e)** Mean gray value of F-actin in lamellipodia. ***p < 0.001; ****p < 0.0001; ns, not significant with 3 ≤ N ≤ 8 replicates per condition.

## Cell migration speed is modulated by matrix chirality

We hypothesized that changes in matrix chirality can modulate the migration speed of epithelial cells. To verify this hypothesis and gain a deeper understanding into the consequences of morphological adaptation to matrix chirality on cell migration, we monitored the movement of individual keratocytes migrating on either COL **(Supplementary Movie 2)**, L- **(Supplementary Movie 3)** and D-CMPs **(Supplementary Movie 4)** with time-lapse microscopy over a 1-hour period. Each epithelial cell was subsequently tracked^30^ to determine the *(x,y)* coordinates of the cell body. We obtained representative trajectories of individual epithelial cells migrating on COL, L- and D-CMPs **(Fig. 4a)** and determined the corresponding mean square displacement (MSD) versus time **(Fig. 4b)**. MSD was lower on D-CMPs and almost similar on COL and L-CMPs, suggesting cells explored larger surface areas on COL and L-CMPs than on D-CMPs. Interestingly, we found no statistical difference of the MSD slope which remained close to 1.8 for all conditions, suggesting that chirality changes do not impact the migration persistence regime of epithelial cells. However, we observed a significant decrease of the cell migration speed on D-CMPs (6.50 ± 2.14 μm/min), while COL and L-CMPs exhibited similar cell migration speed values, with 8.78 ± 3.01 μm/min and 8.64 ± 2.67 μm/min, respectively **(Fig. 4d)**. Altogether, these results demonstrate that epithelial keratocytes can interact and spread on D-surfaces but exhibit a smaller spreading area, a rounded shape and an approximately 25% reduced migration speed.

**Figure 4.**
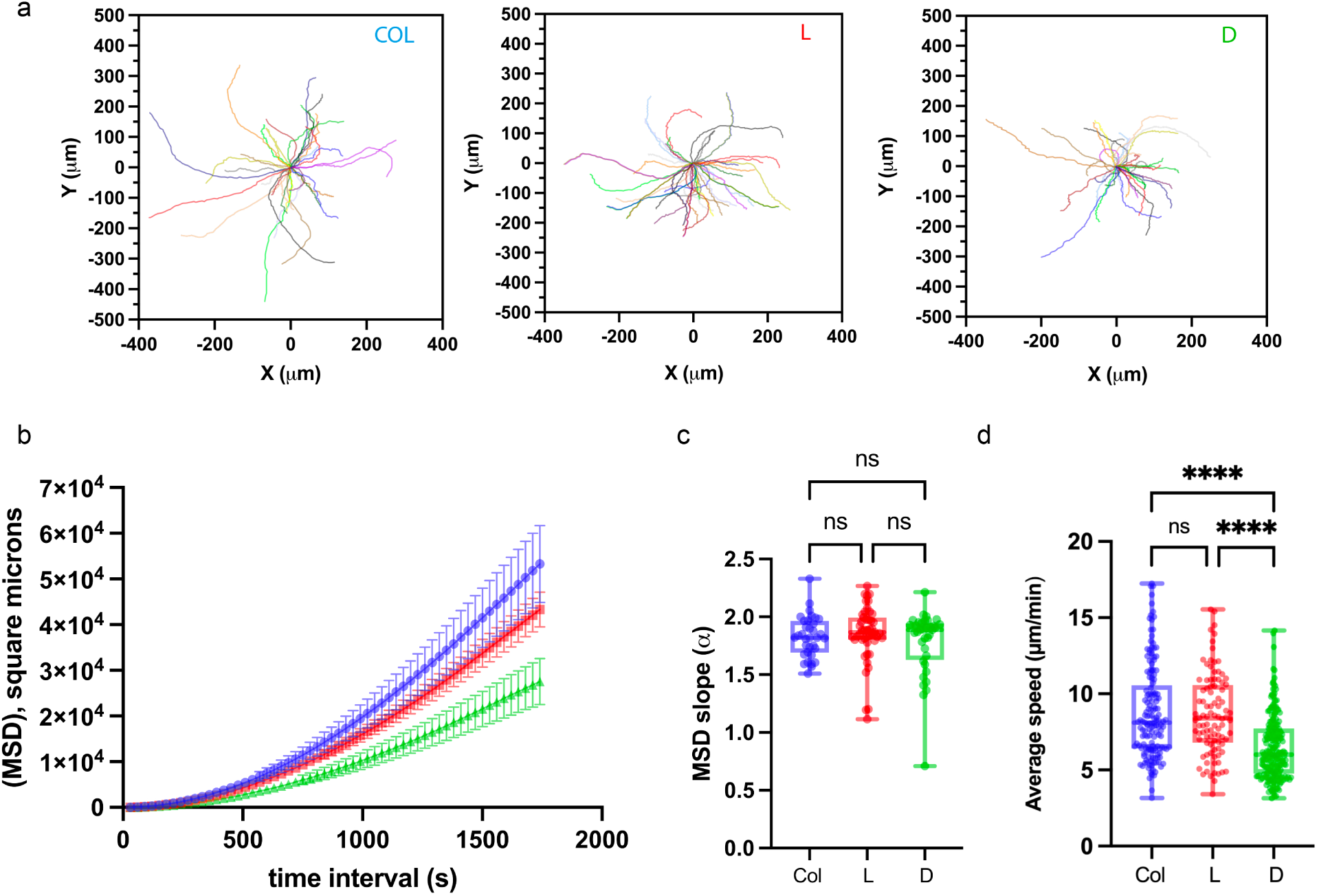
– Matrix chirality modulates cell migration velocity. **(a)** Representative cell trajectories and **(b)** Mean Square Displacement (MSD) versus time interval for COL (blue), L-CMPs (red) and D-CMPs (green). **(c)** MSD slopes on COL (blue, n=34), L-CMPs (red, n=47) and D-CMPs (green, n=36). **(d)** Average cell speed on COL (blue, n=157), L-CMPs (red, n=96) and D-CMPs (green, n=190). ****p < 0.0001; ns, not significant with at least 3 replicates per condition.

## D-chiral surfaces limit the formation of focal adhesions

To comprehensively understand how cells interact with matrices of different chiralities, we focused on cell adhesion sites, crucial for cellular migration^31^. Focal adhesions, situated at the junction between the ECM and the transmembrane integrins^32^, form a dynamic assembly of regulatory and structural proteins capable of transducing external signals and triggering intracellular responses, including integrin activation^33^. Among these proteins, vinculin plays a key role in focal adhesion formation by interacting with talin and actin filaments^34^. Keratocytes cultured on COL, L- and D- CMPs were immunostained for vinculin **(Fig. 5a-c)** by confocal microscopy and fluorescent images were analyzed using a customized pipeline designed for collecting very small adhesions **(Supplementary Fig. 3).** We observed a high number of adhesions on COL (296 ± 79, Fig. 4d), which is not significantly different than for L-CMPs (255 ± 76). However, the number of focal adhesions was significantly lower on D-CMPs (178 ± 52) **(Fig. 5d)**. Our results indicate a lower total area of adhesions on L- and D-CMPs **(Fig. 5e)** than on COL, with a total adhesion area roughly two times smaller on D-CMPs (52 ± 14 µm^2^) than on COL (94 ± 22 µm^2^). Considering our previous results on cell spreading area, we normalized the total adhesion area with the spreading area **(Fig. 5f)**. This yielded a percentage of adhesion, which was comparable on COL (≈16%) and L-CMPs (≈16%) but statistically lower on D-CMPs (≈13%), demonstrating that D-chiral surfaces lead to less specific cell-substrates interactions with epithelial cells.

**Figure 5.**
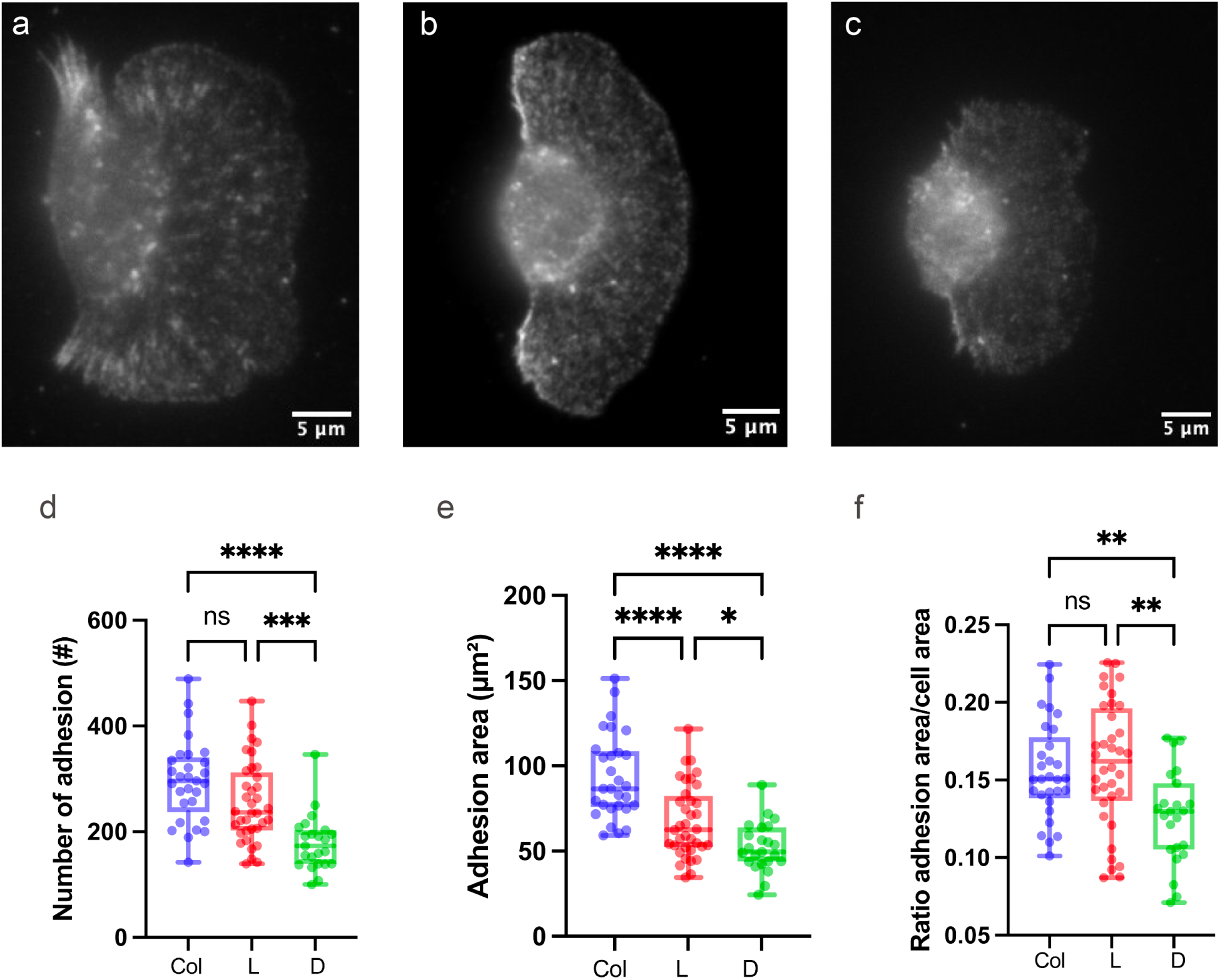
– Modulation of cell adhesions with matrix chirality. Typical fluorescence images of a keratocytes immunostained for vinculin on **(a)** COL, **(b)** L-CMPs, and **(c)** D-CMPs. **(d)** Total number of focal adhesion per cell on COL (blue, n=29), L-CMPs (red, n=39), and D-CMPs (green, n=23). **(e)** Total area of vinculin per cell on COL (blue, n=29), L-CMPs (red, n=39), and D-CMPs (green, n=23). **(f)** Normalized adhesion area on COL in blue (n=29), L-CMPs (n=39), and D-CMPs in green (n=23). All data are shown as mean ± s.d. *p < 0.05; **p < 0.01; ***p < 0.001; ****p < 0.0001; ns, not significant with 3 replicates.

## Integrin inhibition and interactions at the atomic scale

It was shown previously that epithelial keratocytes established close contacts with the matrix through the recruitment of integrins^20,35^. In the subclass of collagen-binding integrins, the molecular recognition process depends on the collagen type^36^. Our results indicated that keratocytes form less specific cell-substrates adhesions on D-chiral surfaces. We hypothesized that the sensitivity of cells to the chirality of the CMPs matrix might result from the interaction between the local conformation of peptides and integrin-type collagen receptors^37^. As it has been shown that α1≈1 is involved in cell migration^38,39^ on collagen type I, we used a α1≈1 blocking antibody to test this hypothesis. After monitoring the migration of control keratocytes for 2 hours using time-lapse microscopy, the α1≈1 blocking antibody was added to the culture medium and the time-lapse was recorded for an additional 2 hours. As shown in **Fig. 6a-c**, the migration speed of treated keratocytes decreased of ≈20% (6.65 ± 3.29 μm/min) on COL (**Supplementary Movie 5)** and ≈25% (6.18 ± 2.31 μm/min) on L-CMPs after α1≈1 inhibition **(Supplementary Movie 6)**, suggesting that α1≈1 is partially involved in the cell migration process. Surprisingly, α1≈1 inhibition on D-CMPs did not affect significantly the migration speed of keratocytes (≈6.70 μm/min, **Supplementary Movie 7**), suggesting subtle differences in interactions between integrins and CMPs.

**Figure 6.**
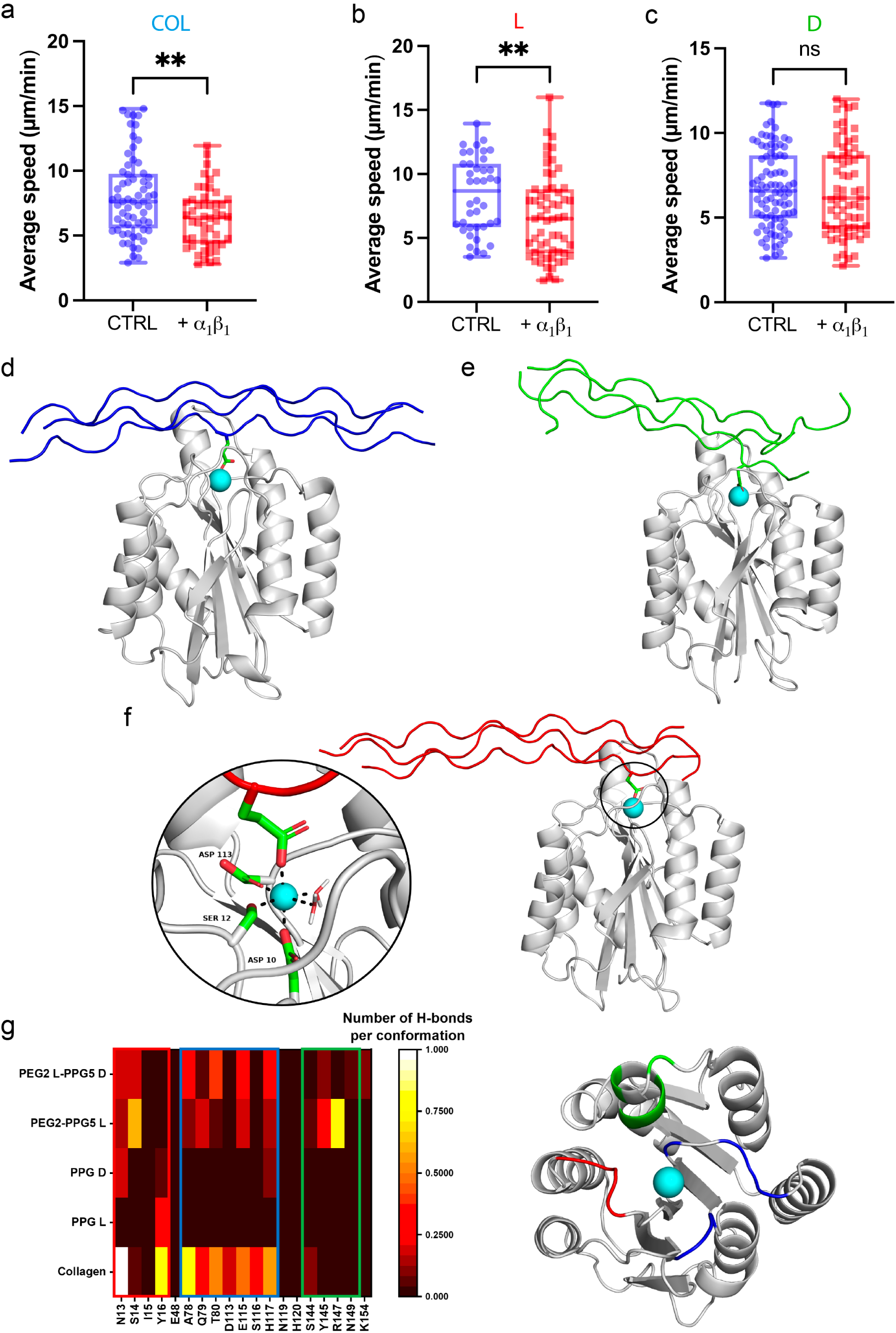
– Inhibition and molecular dynamics simulations of α1β1 integrins. Mean cell migration speed in response to the inhibition of α1β1 **(a)** on COL, **(b)** L-CMPs, and **(c)** D-CMPs matrices. Blue data corresponds to the cell speed before inhibition (n=40 for COL, n=52 L-CMPs, and n=88 for D-CMPs) and red data after inhibition (n=59 for COL, n=45 L-CMPs, and n=67 for D-CMPs). **p < 0.01; and ns, not significant with at least 5 replicates per condition. **(d)** MD snapshot of the collagen triple helix (in blue lines) bound to the I-domain of the α1β1 integrin (in gray). The Mg^2+^ ion is shown as a cyan sphere. **(e)** MD snapshot (last frame) of the simulation of the heterochiral CMP (in green) bound to the I-domain of the α1β1 integrin (in gray). The disordered (PEG)2 part is located on the right of the triple helix, and the Mg^2+^ ion is shown as a cyan sphere. **(f)** MD snapshot (last frame) of the simulation of the homochiral CMP (in red) bound to the I-domain of the α1β1 integrin (in gray). A zoom shows the coordination sphere of the Mg^2+^ ion (in cyan), with three integrin amino acids (ASP 113, SER 12 and ASP 10), two water molecules, and a CMP glutamate residue. **(g)** On the left: heatmap of the intermolecular H-bonds between the peptidic systems (*y*-axis) and the amino acids of the binding site of the α1β1 integrin (*x*-axis). At the crossing of the *x* and *y* axes are displayed rectangles, whose color indicates the frequence of an interaction : the brighter the rectangle, the more frequent the H-bond. Some amino acids are close to each other on the binding site and can be distinguished in three groups depicted with red, blue and green boxes. On the right: top-view of the binding site. The colored parts (in red, blue and green) indicate the positions of the three groups of amino acids, as in the boxes on the heatmap.

To better understand our experimental observations, we conducted atomistic MD simulations to study the intermolecular interactions between the integrin and CMPs of opposite chirality. Collagen-binding integrins all share the presence of an I-domain, responsible for binding^40^. Several studies have highlighted the role of the “GFOGER” sequence in collagen for interactions with the I-domain of α integrins^15^. Considering the high structural similarity of the I-domains of the α1β1 and α2β1 integrins^41^, we reproduced the binding mode of the model collagen with the I-domain of α2β1 for the starting structure of our simulations with α1β1 **(Fig. 6d-g**, see methods for details). As in natural collagen, our CMPs contain glutamate residues, which act as a coordinating unit to a divalent cation located in a metal-ion dependant adhesion site (MIDAS)^42^. In the case of the I-domain, glutamate can coordinate a Mg^2+^ ion in the binding site, thus completing its coordination sphere, as shown in the zoom in **Fig. 6g**. In our simulations on the interaction between each CMP and the I-domain, in all cases the glutamate maintained its interaction with the ion during the whole simulation and the triple helix remained bound to the domain **(Extended Fig. 2a and Supplementary Movies 8-10)**. For comparison, simulations of pure L- or D-(PPG)10 triple helices (without the presence of the PEG triplets) with the I-domain indicated that they were markedly less stabilized, with possible complete unbinding **(Supplementary Movie 11)**. This result shows the crucial role of the glutamate moiety in the interaction with the I-domain, which was also highlighted by others^43^. By coordinating the ion, this interaction anchors the (PPG)10 triple helix in the cavity of the active site, thereby significantly increasing the number of intermolecular H-bonds **(Extended Fig. 2b)**. Interestingly, the homochiral peptide (L-CMP) has slightly more interactions with the domain (2.8 ± 1.6 H-bonds per conformation) than its heterochiral (D-CMP) counterpart (2.2 ± 1.2 H-bonds per conformation). It is in line with the experimental observation that the number of focal adhesions per cell is reduced on D-surfaces.

The model collagen carries out even more H-bonds (5.4 ± 1.5 H-bonds per conformation), thanks to the presence of hydroxyprolines and the GFOGER recognition motif. We have also localized with which amino acids of the binding site are interacting the peptides **(Figure 6g, left)**. The heatmap indicates, on the *x*-axis, all the amino acids which form H-bonds with the collagen during the simulations. Three groups of amino acids can be distinguished, based on their position in the domain **(Figure 6g, right)**. We observe that the CMPs reproduce quite well the interaction pattern of the collagen, although differences arise, essentially in the number of some H-bonds. The homochiral peptide displays two persistent hydrogen bonds, with a serine (S14) and an arginine (R147). In comparison, the heterochiral peptide does not show very persistent interactions, its most frequent H-bond (with tyrosine T80) occurring less than 40 % of the time. The lower number of interactions for the heterochiral peptide likely finds its origin in the chiral mismatch at the junction between the L- and D-amino acids (**Figure 1d**), which disorganizes the triple helix. This junction being precisely located in the binding pocket, the interactions at play between the heterochiral CMP and the I-domain are lower in average compared to those with the homochiral CMP (or collagen). This is also the case for bridging H-bonds, where one amino acid of the triple helix and one amino acid of the I-domain bind to the same water molecule. These interactions are slightly more frequent for the homochiral peptide than for the heterochiral CMP **(Extended Fig. 2c)**. Furthermore, the homochiral L-peptide displays the most persistent bridging interactions, maintained around 90% of the time. It occurs between two glutamate residues, with sometimes two water molecules on the same time. This contact stabilizes the triple helix structure of the homochiral peptide in the (LPLEG)2 region, compared to the heterochiral peptide for which these interactions are weaker and less persistent.

## Conclusions

By design, the two CMPs studied here as cell substrates contain PPII helices of inverse chirality, while their integrin I-domain binding sequences (PEG) possess indentical (L) chirality. For the heterochiral peptide, this creates a chirality mismatch at the junction between L and D amino acids, destabilizing the PPII helix and impairing the molecular recognition of the I-domain compared to the specific interactions with the homochiral sequences. These differences in molecular recognition are evident at the cellular level, particularly when epithelial cells migrate on the CMP substrates. In the case of the homochiral CMP, the migration speed and number of focal adhesions are comparable to those observed on natural collagen substrates. However, introducing a chirality mismatch at the junction between the inverse PPII helix and the PEG recognition motif significantly reduces migration speed and the number of focal adhesions. These findings highlight the extreme sensitivity of cells to chirality mismatches within the matrix, sensed through integrin-mediated molecular recognition. This demonstrates that cells can detect local structural variations at an atomistic level, responding to relatively minor chiral perturbations. Our results lay the groundwork for future studies where cellular responses can be modulated by local chirality, potentially leading to new insights into cell behavior and the development of innovative biomaterials designed to direct cellular functions through precise chiral cues.

## Author contributions

M.S. and S.G. conceived and supervised the project. A.R. developed the substrates, performed cell experiments, tracking, and optical imaging. D.D. and M.S. developed and implemented the molecular simulations. A.R., M.S., and S.G. analyzed the experimental data. D.D. analyzed the theoretical data. M.L. contributed to the discussion and interpretation of the results. The article was written, read, and revised by all authors.

## Acknowledgments.

A.R, M.L., and S.G. acknowledge funding from the University of Mons ‒ UMONS, the FEDER Prostem Research Project no. 1510614 (Wallonia DG06), the F.R.S.-FNRS Epiforce Project no. T.0092.21, the F.R.S.-FNRS Cellsqueezer Project no. J.0061.23, the F.R.S.-FNRS Optopattern Project no. U.NO26.22 and the Interreg MAT(T)ISSE project, which is financially supported by Interreg France-Wallonie-Vlaanderen (Fonds Européen de Développement Régional, FEDER-ERDF). M.S. and D.D. thank acknowledges funding from the University of Mons ‒ UMONS, the Fund for Scientific Research F.R.S.- FNRS and FWO under Excellence of Science (EOS) project PRECISION n°30650939. A.R. is Research Fellow (Aspirant) of the F.R.S.-FNRS, M.L. is Postdoctoral Fellow (Chargé de Recherches) of the F.R.S.- FNRS and M.S. is Research Director of the F.R.S.-FNRS. This work was supported by the Fonds de la Recherche Scientifique – FNRS and by the French Community of Belgium within the framework of a FRIA grant for D.D. Computational resources are provided by the Consortium des Équipements de Calcul Intensif (CÉCI), funded by FNRS (Grant No. U.G.018.18) and Wallonia Region, and Lucia, the Tier-1 supercomputer of the Walloon Region, infrastructure funded by the Walloon Region under the grant agreement n°1910247. We acknowledge EuroCC Belgium, for awarding this project access to the LUMI supercomputer, owned by the EuroHPC Joint Undertaking, hosted by CSC (Finland) and the LUMI consortium.

## Ethical compliance

Primary epithelial cells, keratocytes, harvested from a scale of *hypsophrys nicaraguensis* was done in accordance with European guidelines for animal experimentation and in agreement with the local ethic committee of the University of Mons which reviewed the procedure. A.R. and S.G.

## Data availability

All data are available from the corresponding authors upon request.

## Competing interests

The authors declare no competing interests.

**Correspondence and requests for materials** should be addressed to Mathieu Surin or Sylvain Gabriele.

## Materials and Methods

### Microcontact printing

To create flat micropattern of collagen or collagen-mimetic-peptides, rectangular PDMS stamps were used and created using a silicon wafer. First, the silicon surface was passivated with fluorosilane (tridecafluoro-1,1,2,2-tetrahydrooctyl-1-trichlorosilane) for 30 minutes. Then, the previous PDMS mixture was poured on the silicon wafer and placed 4 hours at 60°C. After the night, the resulting PDMS block was cut in stamps of 1 cm2 approximately. The stamps were washed two times in an ultrasonic bath with a detergent solution (Decon 90, 5%) and then isopropanol solution (70%) for both 15 minutes. Flat stamps were oxidized 7 minutes in an ultraviolet/O3 oven and inked with samples for 1 hours at room temperature. Then, stamps are gently dry by nitrogen flux and pressed onto flat PDMS-coated glass coverslip. Using tweezers, the PDMS stamps were gently removed, and the coverslips were stored in PBS at room temperature in dark room.

### PDMS-coated glass coverslips

Polydimethylsiloxane (PDMS) surfaces were fabricated by mixing the base and the curing agent (Sylgard 184 silicone elastomer kit, Dow Corning Midland, MI) at a specific ratio of 10:1 w/w thoroughly for 5 min and degassed under vaccum for 30 min^44^. The mix was spin-coated at 5,000 r.p.m on 25 mm glass coverslips of 170 µm thick to obtain a thin layer of 25 mm thick. The PDMS layer was then cured for 3 hours at 60°C, flushed with ethanol and exposed to UV illumination for 15 min for sterility. PDMS-coated glass coverslips were stored in the dark at room temperature in a petri dish until use.

### Cell culture

Fish epithelial keratocytes were obtained from the scales of Central American cichlid (*Hypsophrys Nicaraguensis*), as described previously^20,28,29^. Scales were gently taken off the dorsal part of the fish and placed on a glass coverslip previously washed with isopropanol (30%) and dried with nitrogen flux. A drop of 150 ml of fresh Leibovitz’s medium (L15) supplemented with 10% fetal bovine serum (FBS, Capricorn), 1% antibiotic-antimycotic (penicillin/streptomycin, Westburg), 14.2 mM HEPES (Sigma Aldrich), and 30% deionized water was deposited on the scale. A 22-mm glass coverslip sandwiched the scale and epithelial keratocytes were culture at least for 12 hours in dark room with few drops of culture medium around the sample. Individual keratocytes were dissociated by incubating the epithelial tissue with a trypsin solution (1ml per coverslip) during 5 min and resuspended with 4 ml of fresh L15 medium. The suspended cells were centrifugated during 5 minutes at 1200 r.p.m. and then deposited on PDMS-coated glass coverslips. Time-lapse experiments were performed 2 hours after cell seeding. Only polarized cells that actively probed their environment by generating dynamic lamellipodial protrusions were analyzed.

### Molecular dynamics simulations

MD simulations were carried out with the AMBER package^45^. The structures of the I-domain of the α1β1 integrin and of the (PPG)10 triple helix were directly taken from the Protein Data Bank (PDB), with PDB ID: 1qcy^46^ and 1k6f^12^ respectively. To build the collagen-mimetic peptides (PEG)2-(PPG)5, the backbone of the (PPG)10 triple helix was reproduced, and its length and amino acid composition was subsequently adapted with the LEaP module of AMBER. The D-enantiomers were obtained by creating the mirror image of the L-enantiomers using LEaP. The structure of the model collagen of 21 amino acids was extracted from a complex with the I-domain of the α2β1 integrin (PDB ID: 1dzi)^13^. To build the complexes peptide/I-domain, peptides were assembled with the I-domain of the α1β1 integrin by mimicking the binding mode of the model collagen with the I-domain of the α2β1 integrin. This assumption seems reasonable, given the high structural similarity between the I-domains of α1β1 and α2β1^41^. The homochiral and heterochiral collagen-mimetic peptides, the model collagen strand, the L- and D- (PPG)10 triple helices and the I-domain of α1β1 were described with the ff19SB force-field^47^. The complexes peptide/I-domain were solvated in explicit water boxes, using the 4-point OPC water model^48^. NaCl ions were added at a concentration of 0.15 M, using the “SPLIT” method^49^. The simulations started with a geometry optimization performed by molecular mechanics to get a stable starting structure. A first phase served to stabilize the solvent molecules and the Na^+^ and Cl^-^ ions, which underwent 1,000 steps of steepest descent followed by 9,000 steps of conjugated gradient, with restraints on the solute atoms. The second phase of optimization was performed with the same protocol, without any constraints. Next, a heating step of 2 ns was performed in the NVT ensemble. The system was brought to a temperature of 300 K in 1 ns, and was maintained at this temperature for a further 1 ns (and for the rest of the simulation) using a Langevin thermostat, with a collision frequency of 1 ps^-1^. Positional restraints were applied to all solute atoms during heating, with a force-constant of 10 kcal.mol^-1^.Å^-2^. Then, the system was equilibrated during 10 ns in the NPT ensemble at a pressure of 1 bar using a Monte-Carlo barostat, with a pressure relaxation time of 2 ps. Finally, the production phase of 1 µs was launched in the NPT ensemble. Five independent replicas were launched for each complex peptide/I-domain, starting from the same structure optimized by molecular mechanics. A timestep of 2 fs was used with the SHAKE algorithm to constrain bonds involving hydrogen atoms. A cutoff of 12.0 Å was used for non-bonded interactions and the Particle-Mesh Ewald method was used to treat long-range electrostatic interactions. A snapshot was extracted each ns of the production phase for further analyses. The cpptraj module of AMBER and in-house scripts were used to analyze the simulations^50^. RMSF values were computed for each amino acid of the triple helices, after removal of the translational and rotational movements. Hydrogen bonds were detected using geometric criteria: the distance between the acceptor and the donor heavy atom must be < 3.0 Å, and the angle between the donor, the hydrogen atom and the acceptor must be > 135°. The PyMOL 2.5.4 software was used to produce the snapshots and movies^51^.

### Time-lapse imaging and cell tracking

Time-lapse microscopy experiments were carried out on a Nikon Eclipse Ti-E inverted microscope (Nikon C1, Japan) equipped with an automatic stage and a back illuminated sCMOS camera (PCO Edge, Germany). Experiments were performed in Differential Interference Contrast (DIC) mode using a ×20 or ×40 objective to follow the keratocyte migration. Images were taken every 15 seconds and cells were tracked with Matlab (MathWorks) by following their centroids movement at each point. Distance versus time curves were fitted by linear regression to determine the slope corresponding to the mean cell velocity.

### Immunochemistry and confocal microscopy

Fish keratocytes were fixed in 0.05% of glutaraldehyde and permeabilized in 0.1% Triton X-100 for 15 min, followed by three rinses in PBS and a second in 0.2% solution of glutaraldehyde for 10 min. For vinculin staining, cells were permeabilized in a 4% paraformaldehyde solution and 0.1% Triton X-100 for 10 minutes and washed 3 times in PBS. After the fixation and permeabilization, cells were blocked with a solution of 5% FBS in PBS for 30 min and rinsed three times with PBS. Fixed keratocytes were incubated with AlexaFluor 488 phalloidin (Invitrogen, 1:200) to visualize actin filaments and with 4’-6-diamidino-2-phenylindole (DAPI, Invitrogen, 1:200) for the nucleus. For visualizing vinculin containing focal adhesions, cells were first incubated with a primary antibody (anti-vinculin produced in mouse, Sigma-Aldrich, 1:200) for 45 min at 37°C, rinsed three times with PBS, and then incubated with tetramethylrhodamine as secondary antibody (goat anti-mouse, Sigma-Aldrich, 1:200) for 45 min at 37°C. Immunostained keratocytes were mounted in Slow Gold Antifade (Molecular Probes, Invitrogen) and kept at -20°C. Images were collected in confocal and epifluorescence modes with a motorized inverted microscope equipped (Nikon Ti2 A1R HD25, Japan) equipped with a ×60 Plan Apo (NA 1.45, oil immersion) objective and lasers that span the violet (405 and 440 nm), blue (457, 477, and 488 nm), green (514 and 543 nm), yellow-orange (568 and 594 nm), and red (633 and 647 nm) spectral regions. Confocal images were collected with ×100 Plan Apo silicone objective of high numerical aperture (Plan Apochromat Lambda S, Nikon) in galvanometric mode with Z-depth increments of 0.1 mm to capture high resolution images processed by NIS-Elements (Nikon, Advanced Research version 4.5).

### Circular Dichroism spectroscopy

UV-Vis and CD measurements were recorded by using a ChirascanTM Plus CD Spectrometer from Applied Photophysics. The measurements were performed by using 1 mm Suprasil quartz cells (Hellma Analytics) and slide of 1 cm^2^ for the surface characterization. The spectra were recorded between 190 and 300 nm with a bandwidth of 1 nm, time per point 1s. The PBS buffered solution (collagen) and Milli-Q water (peptides) reference spectra were used as baselines and were automatically removed from the CD of the samples. For the thermal analysis, the wavelength was fixed at 220/225 nm (Collagen/Peptides) corresponding to the positive/negative (L/D) peaks. We increased the temperature by 0.5 °C/min, starting at 10°C to 70°C and recorded the signal every 0.25°C. Curves were extracted in csv format and exported in OriginPro v.2021 (OriginLab).

### Scanning Electron Microscopy (SEM)

Fish scales were immediately fixed with gludaraldehyde solution at 0.2% in PBS buffer pH = 7.4 for 24 hours. Fixed scales were rinced in the same PBS buffer and dehydrated in graded ethanol bath. First, the scales were placed in 70% ethanol bath for 30 minutes. Then, a second ethanol bath at 70% was used during 24 hours following by 2 baths of 90% for both 30 minutes. Finally, the scales were placed one hour in a 100% ethanol bath. The scales were dried using a critical-point method by the use of CO2 fluid transition. Then, they were coated with a mix of gold and palladium (40% and 60%, respectively; JFC-1100E metallizer, Jeol) and observed with a Jeol (JSM-7200F) scanning electron microscope.

### Drug treatment

Inhibition of integrins was carried out using an anti-integrin α1β1 antibody (Sigma Aldrich), which was added at 10 µg/ml to the culture media after 2 hours of controlled migration.

### Statistical information

Each experiment was repeated at least three times on each substrate, using different coverslips at different days. Every set of data was tested for normality test using the d’Agostino-Pearson test in Prism 10.0 (GraphPad) that combines skewness and kurtosis tests to test whether the shape of the data distribution was similar to the shape of a normal distribution. For paired comparisons, significances were calculated in Prism 10.0 (GraphPad) with a Student’s t-test (two-tailed, unequal variances) when the distributions proved to be normal. If a data set did not pass the normality tests, the significances were calculated with Mann–Whitney (two-tailed, unequal variances). For multiple comparisons with non-normal distribution, data sets were analyzed with a Kruskal-Wallis test in Prism 10.0 (GraphPad), which is a suitable nonparametric test for comparing multiple independent groups when the data are skewed. When the null hypothesis was not retained (p-value < 0.05), Kruskal-Wallis was corrected with Dunn’s test, which is a nonparametric test with no pairing and multiple comparisons that can be used for both equal and unequal sample sizes. Unless otherwise stated, all data are presented as mean ± standard deviation (s.d.). The confidence interval in all experiments was 95% and as a detailed description of statistical parameters it is included in all figure captions with *p < 0.05, **p < 0.01, ***p < 0.001, ****p<0.0001 and n.s. is not significant.

---

**Supplementary Movie 1 –** MD simulation movie (t=1 µs) of the (PEG)2-(PPG)5 triple helix structure of the CMPs, extracted from their simulation with the integrin. The movie shows the high flexibility of the (PEG)2 region of the heterochiral CMP.

**Supplementary Movie 2 –** Time-lapse sequence in DIC mode of keratocyte migrating on collagen type I. The frame interval is 15 seconds, and the scale bar is 15 μm.F

**Supplementary Movie 3 –** Time-lapse sequence in DIC mode of keratocyte migrating on L-CMPs. The frame interval is 15 seconds, and the scale bar is 15 μm.

**Supplementary Movie 4 –** Time-lapse sequence in DIC mode of keratocyte migrating on D-CMPs. The frame interval is 15 seconds, and the scale bar is 15 μm.

**Supplementary Movie 5 –** Time-lapse sequence in DIC mode of keratocytes migrating on collagen type I after the inhibition of α1≈1. The frame interval is 15 seconds, and the scale bar is 50 μm.

**Supplementary Movie 6 –** Time-lapse sequence in DIC mode of keratocytes migrating on L-peptide matrix after the inhibition of α1≈1. The frame interval is 15 seconds, and the scale bar is 50 μm.

**Supplementary Movie 7 –** Time-lapse sequence in DIC mode of keratocytes migrating on D-peptide matrix after the inhibition of α1≈1. The frame interval is 15 seconds, and the scale bar is 50 μm.

**Supplementary Movie 8 –** MD simulation movie (t=1 µs) of the collagen (in blue) bound to the I-domain of the α1β1 integrin (in gray). The glutamate residue is shown in stick representation, coordinating the Mg^2+^ ion (cyan sphere).

**Supplementary Movie 9 –** MD simulation movie (t=1 µs) of the homochiral CMP (in red) bound to the I- domain of the α1β1 integrin (in gray). The glutamate residue is shown in stick representation, coordinating the Mg^2+^ ion (cyan sphere).

**Supplementary Movie 10 –** MD simulation movie (t=1 µs) of the heterochiral CMP (in green) bound to the I-domain of the α1β1 integrin (in gray). The glutamate residue is shown in stick representation, coordinating the Mg^2+^ ion (cyan sphere).

**Supplementary Movie 11 –** MD simulation movie (t=250 ns) of the pure L-(PPG)10 triple helix (in red) bound to the I-domain of the α1β1 integrin (in gray), showing the disassembly of the peptide. The Mg^2+^ ion is depicted as a cyan sphere.

**Extended Figure 1.**
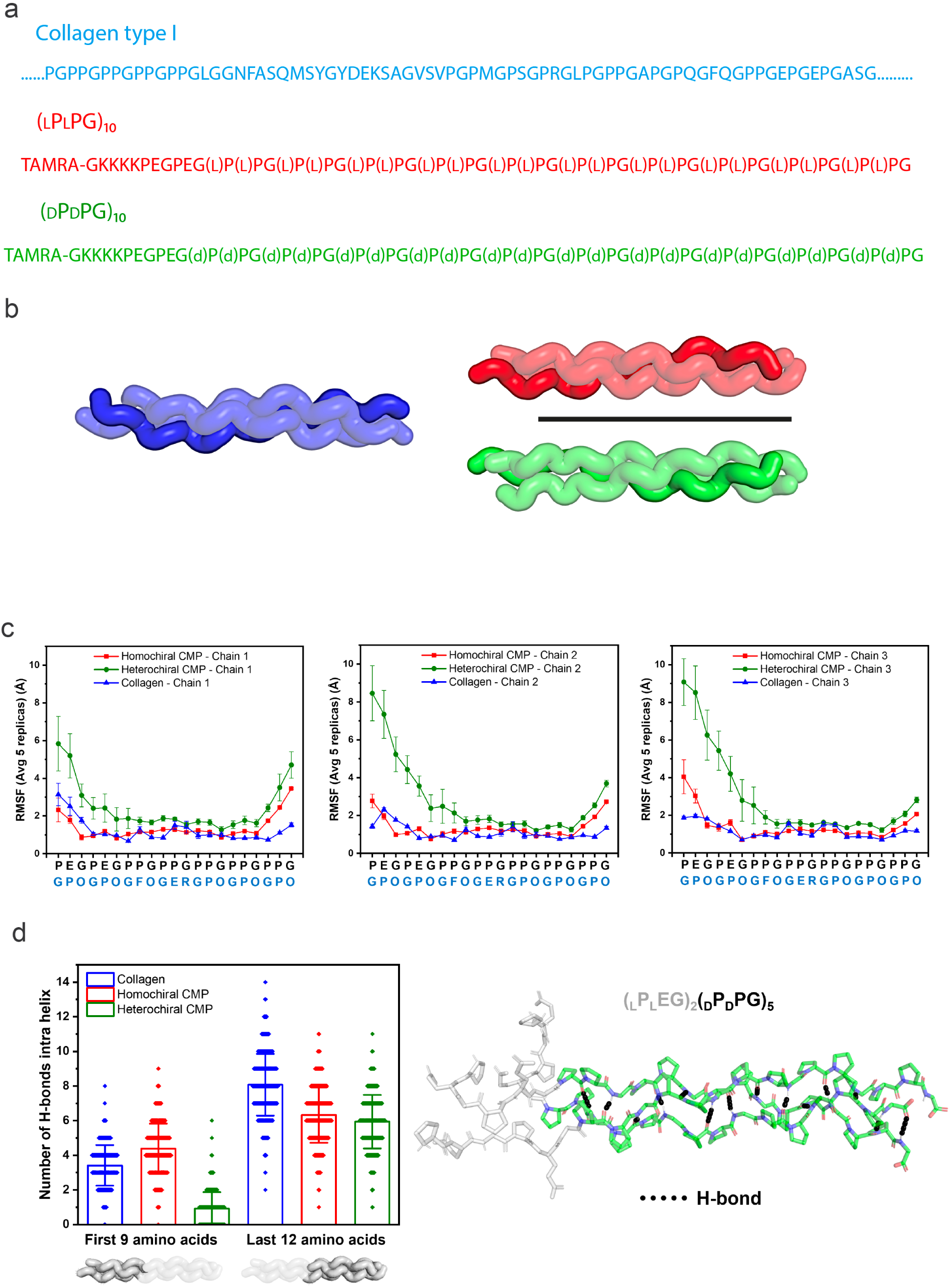
– Characterization of the chiral samples. (**a**) D representations of a short sequence of collagen type I (in blue), the levorotatory [(LPLPG)10, in red], and the dextrorotatory [(DPDPG)10, in green] stereoisomers. (**b)** Visualization of the triple helix resulting from the association of three single chains in the simplified model. (**c**) Root-mean-square fluctuation (RMSF) values for the three peptidic chains forming the triple helix structure (from left to right: first, second, and third chain). Values are averaged over five independent simulations and computed for each CMP amino acid (sequence in black on the x-axis) and collagen (sequence in blue). The higher the RMSF value, the more a residue undergoes position fluctuations. (**d)** Left: Distribution of the number of H-bonds between the first nine and last twelve amino acids of the three peptidic strands inside the triple helix for collagen (blue), homochiral CMP (red), and heterochiral CMP (green). Data is shown as mean ± s.d. Right: MD snapshot of the heterochiral CMP (represented in sticks) showing that H-bonds (displayed as black dots) are maintained in the PPG part, while they are lost in the more disordered (PEG)2 residues (represented as gray sticks).

**Extended Figure 2.**
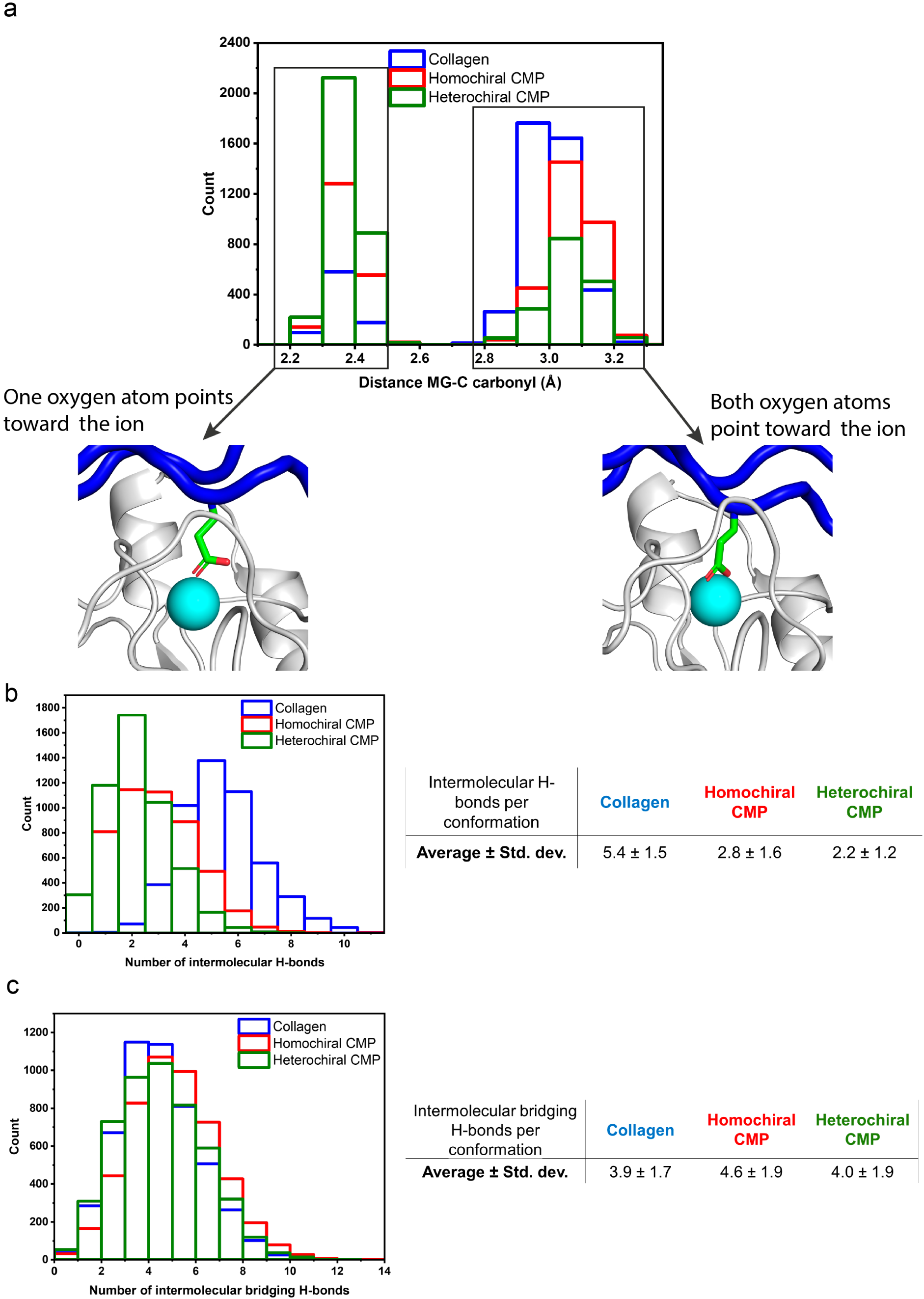
– Study of the interactions between the peptides and the α1≈1 integrin by MD simulations. **(a)** Distribution of the distance between the Mg²⁺ ion and the terminal carbon atom of the coordinated glutamate residue. The distance remained below 3.5 Å for all conformations, indicating that no disassembly occurred. Two distributions appeared, depending on the orientation of the oxygen atoms towards the ion, as shown by MD snapshots. **(b)** Distribution of the number of intermolecular H-bonds per conformation between peptides and the integrin. The average values and standard deviations are provided in the table on the right. **(c)** Distribution of the number of intermolecular bridging H-bonds per conformation between the peptides and the integrin. The average values and standard deviations are provided in the table on the right.

**Supplementary Figure 1.**
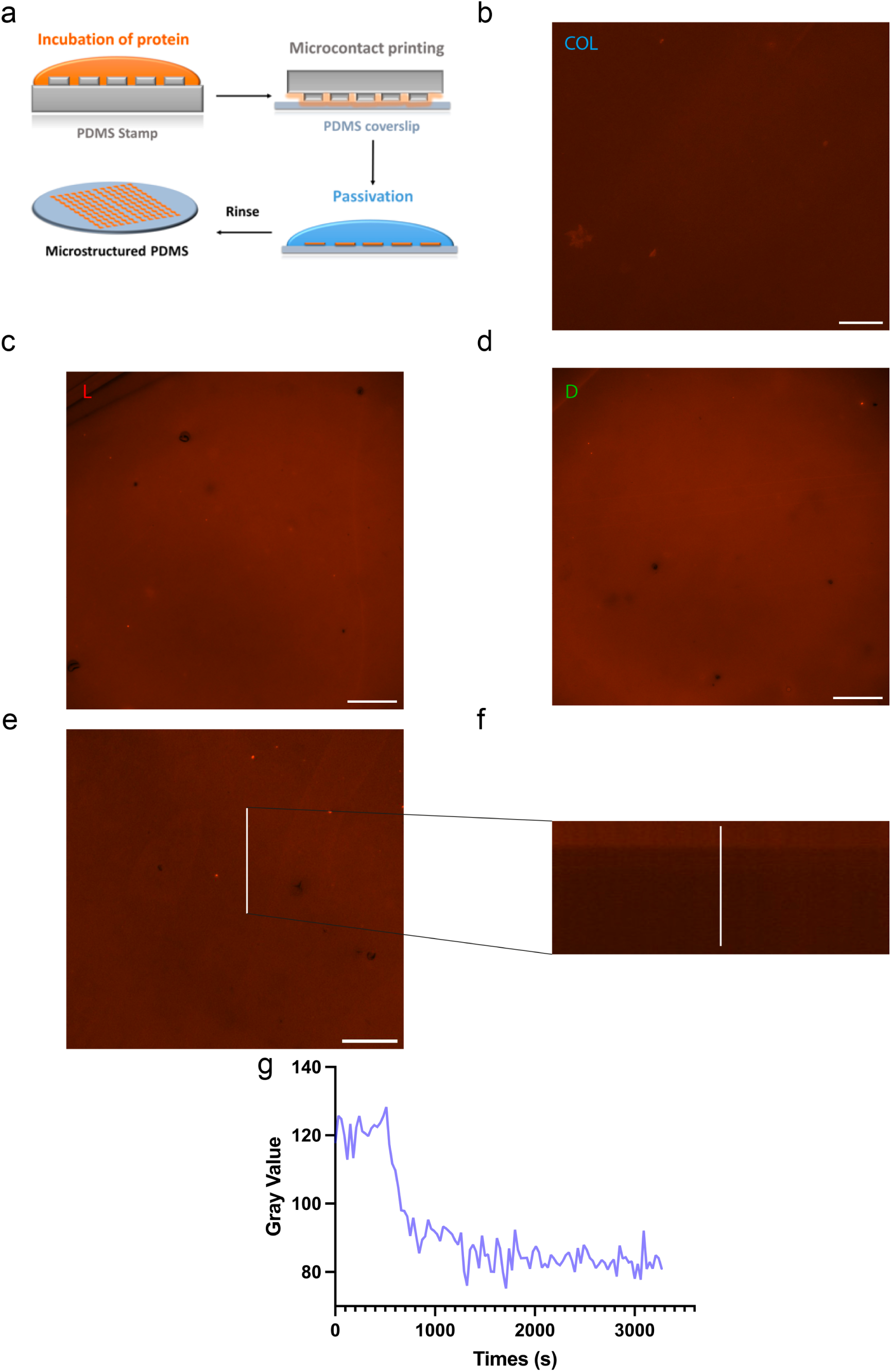
– Stability of the microcontact printing method over time. **(a)** Typical steps of the microcontact printing (μCP) technique used to generate well-controlled functionalized surfaces. Epifluorescent image of **(b)** a collagen-coated surface, **(c)** a L-CMP-coated surface, and **(d)** a D-CMP- coated surface, all generated by μCP. **(e)** Analysis of the L-CMP stability over time based on the examination of **(f)** the mean gray value of a specific section (depicted as a white line) over time. **(g)** Mean gray value over time of the peptide signal with the culture medium added after 300 seconds.. Scale bars correspond to 20 μm.

**Supplementary Figure 2.**
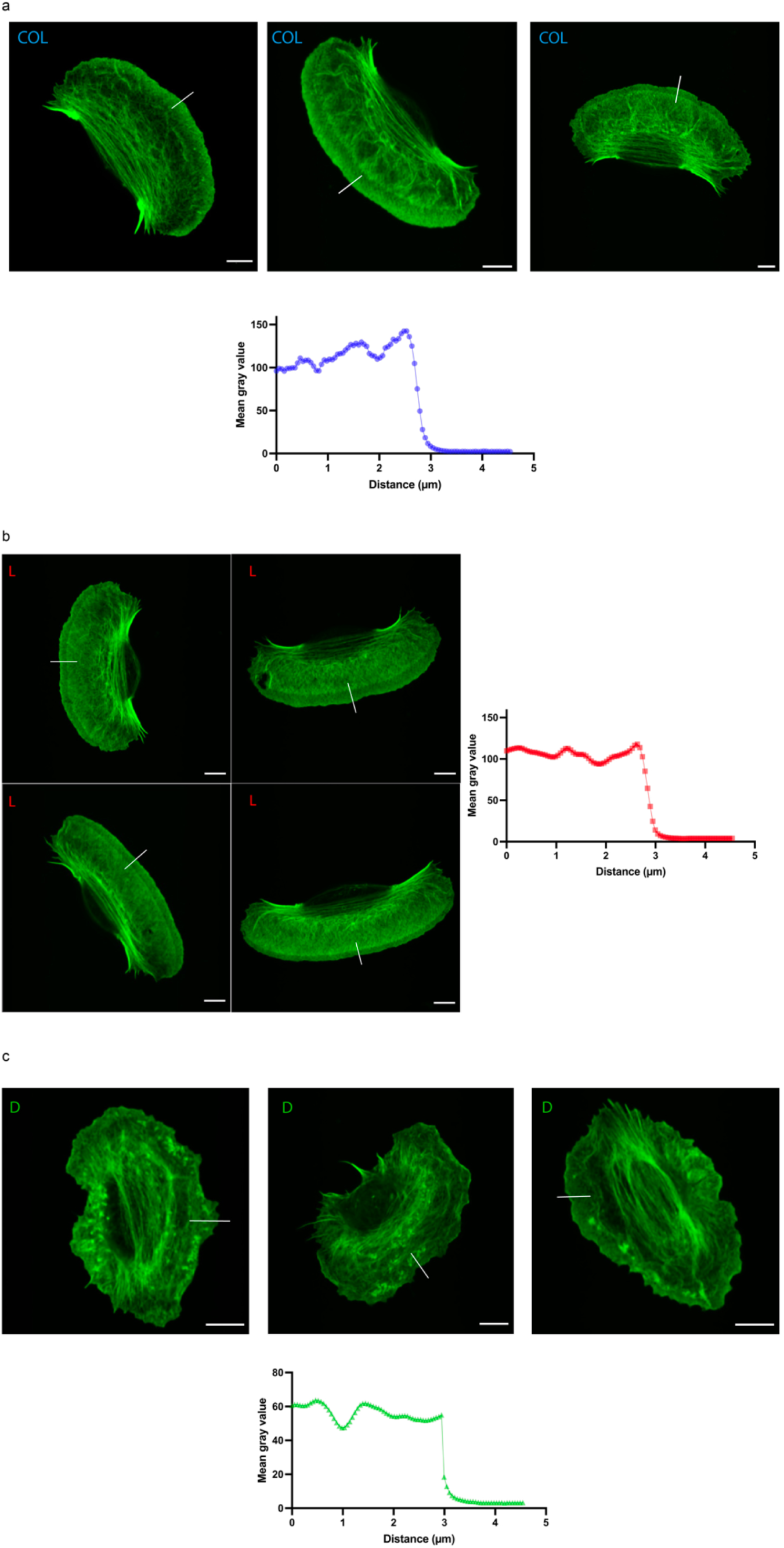
– Confocal analysis of actin distribution at the leading edge of the cell. **(a)** Typical confocal image of keratocyte cells on collagen type I. The white lines indicate the sections where the actin in the lamellipodium was analyzed. The accompanying graph shows the average mean gray value extracted from these sections. **(b)** Typical confocal image of keratocyte cells on L-CMP. The white lines indicate the sections where the actin in the lamellipodium was analyzed. The accompanying graph shows the average mean gray value extracted from these sections. **(c)** Typical confocal image of keratocyte cells on D-CMP. The white lines indicate the sections where the actin in the lamellipodium was analyzed. The graph shows the average mean gray value extracted from these sections. Scale bars are 5 µm.

**Supplementary Figure 3.**
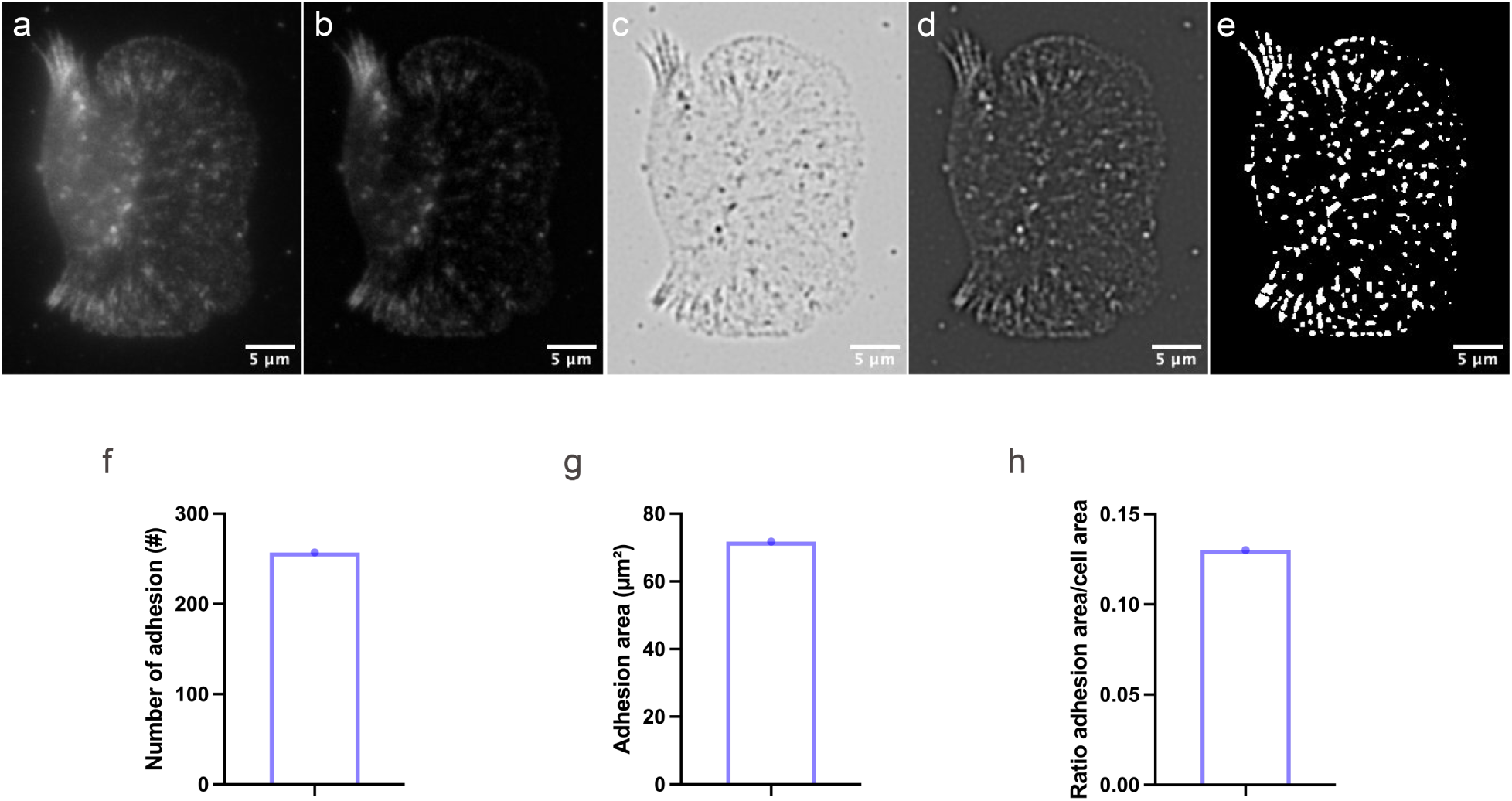
– Pipeline for focal adhesion analysis. **(a)** Epifluorescent image of a keratocyte on collagen type I substrate. The cell is immunostained for vinculin. **(b)** Background subtracted from the previous image to reduce noise. **(c)** Log 3D filter applied to the image to identify small adhesions. **(d)** LUTs inversion of the image. **(e)** Log 3D filter applied to the image to identify small adhesions. **(f)** Total number of focal adhesions for the keratocyte on collagen type I. **(g)** Total area of the adhesions. **(h)** Ratio between the total area of adhesion and the area of the cell.

## References

1. Wood, W. B. Left-right asymmetry in animal development. Annual Review of Cell and Developmental Biology 13, 53–82 (1997).

2. Deng, M., Yu, J. & Blackmond, D. G. Symmetry breaking and chiral amplification in prebiotic ligation reactions. Nature 626, 1019–1024 (2024).

3. Hanein, D., Geiger, B. & Addadi, L. Differential Adhesion of Cells to Enantiomorphous Crystal Surfaces. Science 263, 1413–1416 (1994).

4. Yao, X., Hu, Y., Cao, B., Peng, R. & Ding, J. Effects of surface molecular chirality on adhesion and differentiation of stem cells. Biomaterials 34, 9001–9009 (2013).

5. Männer, J. On the form problem of embryonic heart loops, its geometrical solutions, and a new biophysical concept of cardiac looping. Annals of Anatomy - Anatomischer Anzeiger 195, 312–323 (2013).

6. Lebreton, G. et al. Molecular to organismal chirality is induced by the conserved myosin 1D. Science 362, 949–952 (2018).

7. Tee, Y. H. et al. Cellular chirality arising from the self-organization of the actin cytoskeleton. Nat Cell Biol 17, 445–457 (2015).

8 Chin, A. S. et al. Epithelial Cell Chirality Revealed by Three-Dimensional Spontaneous Rotation. Proc. Natl. Acad. Sci. U.S.A. 115, 12188–12193 (2018).

9. Blackmond, D. G. The origin of biological homochirality. Cold Spring Harb Perspect Biol 2, a002147–a002147 (2010).

10. Koohestani, F. et al. Extracellular matrix collagen alters cell proliferation and cell cycle progression of human uterine leiomyoma smooth muscle cells. PLoS One 8, e75844–e75844 (2013).

11. Smith, L. R., Cho, S. & Discher, D. E. Stem Cell Differentiation is Regulated by Extracellular Matrix Mechanics. Physiology 33, 16–25 (2018).

12. Berisio, R., Vitagliano, L., Mazzarella, L. & Zagari, A. Crystal structure of the collagen triple helix model [(Pro-Pro-Gly)(10)](3). Protein Sci 11, 262–270 (2002).

13. Emsley, J., Knight, C. G., Farndale, R. W., Barnes, M. J. & Liddington, R. C. Structural Basis of Collagen Recognition by Integrin α2β1. Cell 101, 47–56 (2000).

14. Yonath, A. & Traub, W. Polymers of tripeptides as collagen models: IV. Structure analysis of poly(l-prolyl-glycyl-l-proline). Journal of Molecular Biology 43, 461–477 (1969).

15. Knight, C. G. et al. The Collagen-binding A-domains of Integrins α1β1 and α2β1Recognize the Same Specific Amino Acid Sequence, GFOGER, in Native (Triple-helical) Collagens*. Journal of Biological Chemistry 275, 35–40 (2000).

16. O’Leary, L. E. R., Fallas, J. A., Bakota, E. L., Kang, M. K. & Hartgerink, J. D. Multi-hierarchical self-assembly of a collagen mimetic peptide from triple helix to nanofibre and hydrogel. Nature Chemistry 3, 821–828 (2011).

17 Greenfield, N. J. Using circular dichroism spectra to estimate protein secondary structure. 15.

18. Hunter, C. A. & Anderson, H. L. What is Cooperativity? Angewandte Chemie International Edition 48, 7488–7499 (2009).

19. Chen, J., Peng, Q., Peng, X., Zhang, H. & Zeng, H. Probing and Manipulating Noncovalent Interactions in Functional Polymeric Systems. Chem. Rev. 122, 14594–14678 (2022).

20. Riaz, M., Versaevel, M., Mohammed, D., Glinel, K. & Gabriele, S. Persistence of fan-shaped keratocytes is a matrix-rigidity-dependent mechanism that requires α5β1 integrin engagement. Sci Rep 6, 34141 (2016).

21. Wang, Y. L. Exchange of actin subunits at the leading edge of living fibroblasts: possible role of treadmilling. Journal of Cell Biology 101, 597–602 (1985).

22. Theriot, J. A. & Mitchison, T. J. Actin microfilament dynamics in locomoting cells. Nature 352, 126–131 (1991).

23. Mogilner, A. & Oster, G. Cell motility driven by actin polymerization. Biophysical Journal 71, 3030–3045 (1996).

24. Keren, K. et al. Mechanism of shape determination in motile cells. Nature 453, 475–480 (2008).

25. Lin, C.-H. & Forscher, P. Growth cone advance is inversely proportional to retrograde F-actin flow. Neuron 14, 763–771 (1995).

26. Zhu, D. et al. Structure and Mechanical Performance of a “Modern” Fish Scale. Advanced Engineering Materials 14, B185–B194 (2012).

27. Versaevel, M., Grevesse, T. & Gabriele, S. Spatial coordination between cell and nuclear shape within micropatterned endothelial cells. Nature Communications 3, 671 (2012).

28. Vercruysse, E. et al. Geometry-driven migration efficiency of autonomous epithelial cell clusters. Nature Physics (2024) doi:10.1038/s41567-024-02532-x.

29. Mohammed, D. et al. Substrate area confinement is a key determinant of cell velocity in collective migration. Nat. Phys. 15, 858–866 (2019).

30. Piccinini, F., Kiss, A. & Horvath, P. CellTracker (not only) for dummies. Bioinformatics 32, 955– 957 (2016).

31. DeMali, K. A., Wennerberg, K. & Burridge, K. Integrin signaling to the actin cytoskeleton. Current Opinion in Cell Biology 15, 572–582 (2003).

32. Mitra, S. K., Hanson, D. A. & Schlaepfer, D. D. Focal adhesion kinase: in command and control of cell motility. Nature Reviews Molecular Cell Biology 6, 56–68 (2005).

33. Ridley, A. J. et al. Cell Migration: Integrating Signals from Front to Back. Science 302, 1704– 1709 (2003).

34. Humphries, J. D. et al. Vinculin controls focal adhesion formation by direct interactions with talin and actin. J Cell Biol 179, 1043–1057 (2007).

35. Valiente-Alandi, I., Schafer, A. E. & Blaxall, B. C. Extracellular matrix-mediated cellular communication in the heart. J Mol Cell Cardiol 91, 228–237 (2016).

36. Tulla, M. et al. Selective Binding of Collagen Subtypes by Integrin α1I, α2I, and α10I Domains*. Journal of Biological Chemistry 276, 48206–48212 (2001).

37. Peterson, C. M., Helterbrand, M. R. & Hartgerink, J. D. Covalent Capture of a Collagen Mimetic Peptide with an Integrin-Binding Motif. Biomacromolecules 23, 2396–2403 (2022).

38. Pozzi, A., Wary, K. K., Giancotti, F. G. & Gardner, H. A. Integrin α1β1 Mediates a Unique Collagen-dependent Proliferation Pathway In Vivo. Journal of Cell Biology 142, 587–594 (1998).

39. Moir, L. M., Black, J. L. & Krymskaya, V. P. TSC2 modulates cell adhesion and migration via integrin-α1β1. American Journal of Physiology-Lung Cellular and Molecular Physiology 303, L703– L710 (2012).

40. Barczyk, M., Carracedo, S. & Gullberg, D. Integrins. Cell and Tissue Research 339, 269–280 (2010).

41. Nolte, M. et al. Crystal structure of the α1β1 integrin I-domain: insights into integrin I-domain function. FEBS Letters 452, 379–385 (1999).

42. Campbell, I. D. & Humphries, M. J. Integrin Structure, Activation, and Interactions. Cold Spring Harbor Perspectives in Biology 3, a004994–a004994 (2011).

43. Bella, J. & Berman, H. M. Integrin–collagen complex: a metal–glutamate handshake. Structure 8, R121–R126 (2000).

44. Hamieh, M. et al. Influence of Substrate Properties on the Dewetting Dynamics of Viscoelastic Polymer Films. The Journal of Adhesion 83, 367–381 (2007).

45. Case, D. A. et al. The Amber biomolecular simulation programs. Journal of Computational Chemistry 26, 1668–1688 (2005).

46. Nymalm, Y. et al. Jararhagin-derived RKKH Peptides Induce Structural Changes in α1I Domain of Human Integrin α1β1*. Journal of Biological Chemistry 279, 7962–7970 (2004).

47. Tian, C. et al. ff19SB: Amino-Acid-Specific Protein Backbone Parameters Trained against Quantum Mechanics Energy Surfaces in Solution. J. Chem. Theory Comput. 16, 528–552 (2020).

48. Izadi, S., Anandakrishnan, R. & Onufriev, A. V. Building Water Models: A Different Approach. J. Phys. Chem. Lett. 5, 3863–3871 (2014).

49. Machado, M. R. & Pantano, S. Split the Charge Difference in Two! A Rule of Thumb for Adding Proper Amounts of Ions in MD Simulations. J. Chem. Theory Comput. 16, 1367–1372 (2020).

50. Roe, D. R. & Cheatham, T. E. I. PTRAJ and CPPTRAJ: Software for Processing and Analysis of Molecular Dynamics Trajectory Data. J. Chem. Theory Comput. 9, 3084–3095 (2013).

51. The PyMOL Molecular Graphics System, Version 2.5.4 Schrödinger, LLC.

